# Stationary phloem proteins and their effects on viruses, aphids, and cyst nematodes in Arabidopsis

**DOI:** 10.1101/2025.09.10.675357

**Authors:** Janna Damen, Jaap-Jan Willig, Nyasha Chikwature, Jessica de Haan, Jordi N. Kenter, Marcel Dicke, Monique M. van Oers, Geert Smant, Emilyn E. Matsumura, Karen J. Kloth

## Abstract

The phloem is a specialized tissue that facilitates systemic transport of carbohydrates and signal molecules, making it a common target for viruses, phloem-feeding insects, and cyst nematodes. In Arabidopsis, the stationary phloem-associated SIEVE ELEMENT-LINING CHAPERONE1 (SLI1) and RESTRICTED TEV MOVEMENT (RTM) proteins restrict insect phloem-feeding and potyviruses systemic transport, respectively. However, their broader roles in plant-attacker interactions remain largely unexplored. We investigated the roles of SLI1, RTM1, RTM2, and RTM3 in tobacco etch virus (TEV) infection, as well as in *Myzus persicae* and *Heterodera schachtii* infestations using Arabidopsis mutants. Systemic TEV movement was quantified, aphid behaviour and reproduction were assessed, and cyst nematode infection was monitored. SLI1 did not restrict TEV systemic movement, RTM3 reduced *M. persicae* reproduction without altering feeding behaviour, SLI1, RTM2, and RTM3 supported *H. schachtii* infection and feeding site expansion, and confocal images indicated a possible role of the proteins in *de novo* synthesis of sieve tubes around syncytia. These findings reveal that stationary phloem proteins exert dual and target-specific effects, limiting some attackers while inadvertently facilitating others. This highlights the complexity of phloem-based immunity and underscores the need to unravel its underlying mechanisms to develop strategies to reduce multiple pest and pathogen burdens simultaneously.

**Highlight:** Stationary phloem proteins SLI1, RTM1, RTM2, and RTM3 exert dual and target-specific effects against tobacco etch virus, *Myzus persicae*, and *Heterodera schachtii*.

## Introduction

The phloem is a vital plant tissue. It facilitates the systemic (long-distance) transport of sugars, lipids, amino acids, nucleic acids, proteins, and signal molecules, all essential for plant growth, development, and responses to the environment (Turgeon and Wolf, 2009; De Schepper *et al*., 2013; Kehr *et al*., 2022). However, this highly specialized transport tissue is also commonly targeted by many plant-interacting organisms. For instance, the phloem is an excellent systemic transport medium for pathogens, such as viruses, and its contents make it a carbohydrate-rich food source for both pathogens and pests such as aphids and nematodes (Haider *et al*., 2024). Because the phloem is deeply embedded by multiple cell layers, pathogens have evolved various strategies to access this tissue for their own benefit (Leisner and Turgeon, 1993; Böckenhoff *et al*., 1996; Will and van Bel, 2006). For instance, once a plant virus establishes replication in the initially infected cell, their viral movement proteins modify plasmodesmata to facilitate intercellular movement. After crossing several borders, viruses eventually reach the phloem, where they can spread systemically throughout the entire plant (Schoelz *et al*., 2011; Wang, 2021; Xue *et al*., 2023). Other examples include plant-parasitic cyst nematodes, which induce *de novo* formation of sieve tubes to allocate valuable plant assimilates to their feeding site, and aphids, which directly target the sieve tube for feeding with their needle-like organ (Wyss, 1992; Will and van Bel, 2006).

The phloem is a highly specialized tissue which consists of different cell types. Phloem sap transport is mediated by sieve tubes which consist of sieve elements (SEs) that are connected by callose-rich sieve plates (Knoblauch and van Bel, 1998; Van Bel, 2003; Kalmbach and Helariutta, 2019). Sap flow can be regulated through callose deposition at these sieve plates (i.e., slow down, or a complete halt). The living SE cells are mostly empty to facilitate phloem transport, which is a result of their cell differentiation process wherein the majority of organelles are lost, as well as their transcriptional- and translational machinery. As vital organelles are absent in mature SEs, essential proteins, RNAs, metabolites, and other molecules are provided by their neighbouring companion cells via specialized plasmodesmata pore units, that allow for bidirectional transport of these compounds between CCs and SEs (van Bel and Knoblauch, 2000). Because SEs contain only a minimal set of cellular components after maturation, including endoplasmic reticulum and mitochondria, sieve-tube responses to biotic stressors are thought to be primarily regulated post-translationally, and influenced by surrounding cells. The proteins that localize to the mature SE can either traffic along the sap translocation stream or remain stationary through association with membranes of residual organelles (Sanden and Schulz, 2021). However, little is known about the role of stationary phloem proteins in pathogen infection.

To counteract pathogen and pest attacks, the phloem must also harbour specific defence mechanisms against these intruders. However, the phloem is one of the least understood plant tissues and details of its defence mechanisms remain largely unknown. One of the known phloem-associated resistance mechanisms against viruses is mediated by the *RESTRICTED TEV MOVEMENT* (*RTM*) gene complex proteins RTM1, RTM2, and RTM3, which are stationary phloem proteins that belong to the jacalin-repeat, small heat-shock-like (hsp20-like), and meprin and tumour necrosis factor receptor-associated factor (TRAF)-domain containing protein families, respectively. Each gene in the *RTM* gene complex confers dominant resistance by restricting the systemic transport of some potyviruses including tobacco etch virus (TEV), while still supporting virus replication and movement in the local infected leaf (Mahajan *et al*., 1998; Whitham *et al*., 2000; Chisholm *et al*., 2001; Cosson *et al*., 2010). Interestingly, another phloem-associated stationary hsp20-like protein is SIEVE ELEMENT-LINING CHAPERONE1 (SLI1) which has been identified as a resistance factor against *Myzus persicae* (the green peach aphid) and other phloem-feeding insects. SLI1 reduced phloem-feeding activities, increased salivation in the phloem, and reduced the excretion rate of honeydew droplets by aphids (Kloth *et al*., 2017, 2021). SLI1 and RTM2 are closely related hsp20-like proteins in the UAPVI clade, characterized by a predicted N-terminal alpha-crystallin domain and a C-terminal transmembrane domain (Bondino *et al*., 2012). However, the precise mode of action and the involvement of additional phloem-associated proteins underlying *RTM-* and *SLI1-*mediated resistance remain open questions (Kloth and Kormelink, 2020). On the other hand, plant-parasitic cyst nematodes establish long-term interactions by modifying root cells into specialized feeding structures known as syncytia (Wyss and Zunke, 1986; Wyss, 1992). These structures integrate with the plant’s vascular system, ensuring a continuous flow of nutrients to the nematode throughout its life cycle (Hoth *et al*., 2005, 2008). Although some phloem-related genes show differential expression in nematode-infected roots, the role of stationary phloem proteins in determining resistance or susceptibility to cyst nematodes remains largely unexplored.

Together, these examples highlight how the phloem, despite being a critical transport tissue, is also a major battleground for pest and pathogen interactions with plants. Understanding how stationary phloem proteins contribute to resistance or susceptibility will provide new insights into plant defence strategies and may uncover novel mechanisms to counteract a broad range of pathogens. In this study, we investigated the role of the stationary phloem proteins SLI1, RTM1, RTM2, and RTM3 in pest and pathogen interactions with plants. We hypothesized that the two hsp20-like proteins, SLI1 and RTM2, would have similar functions against both potyviruses and aphids, as well as that these four stationary phloem proteins that are expressed in both root and shoot would negatively affect cyst nematodes (Chisholm *et al*., 2001; Kloth *et al*., 2017; Sanden and Schulz, 2022). To test this, we exposed Arabidopsis T-DNA insertion mutants for these genes to TEV, *M. persicae*, or the beet cyst nematode *Heterodera schachtii* and measured systemic virus movement, aphid feeding and reproduction, and the development of both nematodes and their feeding sites.

## Materials and methods

### Plants and growth conditions

*Arabidopsis thaliana* Col-0 accession (CS60000), homozygous T-DNA insertion lines *sli1-1* (SALK_027475) and *sli1-2* (SAIL_1269_C01), and a *pSLI1:EYFP:SLI1* reporter line in the Col-0 background were available from Kloth et al. (2017). Additionally, the homozygous T-DNA insertion lines *rtm1-2* (WISCDSLOX453-456K7), *rtm1-3* (WISCDSLOX506H02), *rtm2-1* (SALK_010448), *rtm3-3* (GK-801D05-023411), and *rtm3-4* (SAIL_790_D11) all in the Col-0 background, *rtm2-4* (SAIL_40_C07) in the Col-3 background, and the Col-3 accession (CS8846) were obtained from the European Arabidopsis Stock Centre (NASC), followed by selection of homozygous lines by PCR, using primers presented in Supplementary Table S1. Loss-of-function in the T-DNA insertion lines *sli1-1, sli1-2*, and *rtm2-1* was already confirmed by previous studies (Decroocq *et al*., 2006; Kloth *et al*., 2017). The other *rtm* T-DNA insertion lines were tested with reverse transcription quantitative PCR (RT-qPCR) to confirm aberrant expression of the gene of interest (Supplementary Table S2, Fig. S1, RT-qPCR procedure as described below). For the virus and aphid experiments, Arabidopsis seeds were stratified for three days on moist filter paper at 4°C. Seeds were sown into 5-cm diameter pots containing Arabidopsis potting soil (Lentse Potgrond), followed by an Entonem treatment (Koppert Biological Systems, Berkel en Rodenrijs, the Netherlands) to control sciarid flies. Plants were grown in a climate room at 26 ± 1°C during the day and 23 ± 0.5°C at night, with a 16-h-light : 8-h-dark (16:8 L:D) photoperiod, and 50-70% relative humidity, because 26°C induces *SLI1* expression (Kloth *et al*., 2017).

### Validation of Arabidopsis T-DNA insertion lines by RT-qPCR

To validate whether Arabidopsis T-DNA insertion lines had aberrant expression of the target gene, one leaf per sample was collected and flash-frozen in liquid nitrogen for further processing. Tissues were homogenized using a pellet pestle and RNA was isolated with the ISOLATE II RNA Plant Kit (Meridian Bioscience, OH, USA). The RNA was further treated with RQ1 RNase-Free DNase (Promega, WI, USA) according to the manufacturer’s protocol. RNA quantity was measured by using the DeNovix DS-11 FX Spectrophotometer/Fluorometer. Subsequently, DNA-free RNA (∼1000 ng/μl) was converted to cDNA with the SensiFAST™ cDNA Synthesis Kit (Meridian Bioscience, OH, USA). As the genes of interest are only expressed in (mature) phloem cells, we expected low RNA yield and, therefore, diluted the RNA only two-fold with nuclease-free water. RT-qPCR was prepared with the SensiFAST™ SYBR® No-ROX Kit (Meridian Bioscience, OH, USA) according to the manufacturer’s protocol and performed on a Bio-rad CFX96 Touch™ Real-Time PCR Detection System. *SLI1, RTM1, RTM2*, and *RTM3* transcript abundance was analysed using gene specific primers, where *ACT2 (ACTIN 2*, At3g18780) was used as a reference gene (Supplementary Table S2). The average of two technical replicates per gene was used to calculate the ΔCq values. Relative expression was compared between each mutant plant line and its wild type with a Mann Whitney U test and displayed with support of the package “emmeans” (Lenth *et al*., 2025) and “tidyverse” (Wickham *et al*., 2019) in R Software v4.4.1 (R Core Team, 2024).

### Virus infection assay

Tobacco etch virus isolate PV 308 was obtained from the business unit Biointeractions & Plant Health of Wageningen Research (Wageningen, the Netherlands), propagated in *Nicotiana benthamiana*, and stored at -70°C. Infective plant tissue was ground (1 : 1, w/v) in phosphate buffer (0.01 M phosphate buffer, pH 7.0, 0.02 M sodium sulphite) immediately before inoculation. Ten 14-day-old Arabidopsis Col-0, *rtm2-1, sli1-1*, and *sli1-2* plants were mechanically rub-inoculated using carborundum powder (Hull, 2009). At 17 days post-inoculation (DPI), inflorescence tissues were sampled by placing the inflorescences into a 1.5 mL Eppendorf tube and isolating the tissue from the plant by closing the lid, thus preventing cross-contamination. The samples were directly flash-frozen in liquid nitrogen and stored at -70°C until processing.

### Virus detection by RT-qPCR

For RNA isolation, Arabidopsis inflorescence tissue was ground in liquid nitrogen using a pellet pestle. RNA was extracted using the FavorPrep™ Plant Total RNA Kit (Favorgen Biotech Corp, Ping Tung, Taiwan) according to manufacturer’s protocol, including the DNaseI (New England Biolabs, MA, USA) treatment step. Subsequently, cDNA was synthesized from the isolated RNA (150 ng/μL) using M-MuLV Reverse Transcriptase (New England Biolabs, MA, USA) and the Oligo(dT)_18_ reverse primer according to manufacturer’s protocol. TEV transcript abundance was analysed through RT-qPCR using specific primers targeting the TEV cylindrical inclusion helicase region (Supplementary Table S3). *F-BOX* (galactose oxidase/kelch repeat superfamily protein, At5g15710) and *MON1* (*MONENSIN SENSITIVITY1*, At2g28390) were used as reference genes as they are stable in virus-infected Arabidopsis (Lilly *et al*., 2011). Primer efficiencies were determined by a two-fold dilution series (0 to 128 times) using pooled cDNA from ten *rtm2-1* TEV inoculated plants, with efficiencies ranging between 98.5% and 103.3% (Supplementary Table S3). qPCR was performed in a Bio-Rad CFX96 Touch Real-Time PCR Detection System using 1 μL of 16 fold diluted cDNA in 10 μL reactions of the PowerTrack™ SYBR Green Master Mix (A46109, Thermo Fisher Scientific, Vilnius, Lithuania), following the standard cycling conditions as described by the manufacturer, followed by a melt curve. Each sample was prepared as two technical replicates, and each plate included a control for each primer pair containing MilliQ instead of cDNA. As TEV negative samples lack PCR amplification with TEV specific primers, no Cq values are obtained for these samples, and therefore their Cq values were set to 40 in order to perform calculations. ΔCq values were calculated by subtracting the geometric mean of the Cq values of the two reference genes from the average Cq value for TEV. The ΔCq values were normalized using an exponential decay function defined as 2^-ΔCq^. Significant differences between biological groups were determined with the Kruskal-Wallis test. Post-hoc pairwise comparison analyses were conducted using Dunn’s test, followed by a correction for multiple testing according to the Benjamini-Hochberg method. Data analysis and visualization was performed with R Software v4.4.1 (R Core Team, 2024) using the packages “tidyverse”, and “dunn.test” (Wickham *et al*., 2019; Dinno, 2024).

### Aphid reproduction assay

Green peach aphids, *M. persicae*, were reared on radish plants, *Raphanus sativus* at 23 ± 1°C, with a 16:8 L:D photoperiod, and 50% to 70% RH. The aphid reproduction assay on Arabidopsis was performed in a climate chamber under the same plant-rearing conditions mentioned under “Plants and growth conditions”. Each Arabidopsis T-DNA insertion line and its respective wild-type plant line was assessed in separate experiments, conducted in one or more batches on consecutive days (*rtm1-2:* 3, *rtm1-3*: 2, *rtm2-1*: 3, *rtm2-1*: 2, *rtm3-3*: 1, and *rtm3-4*: two batches), with both the mutant and wild type plants included on each experimental day. The day before infestation, adult aphids were placed on a *R. sativus* leaf in a closed glass Petri dish (Ø 10 cm) to produce offspring. On the day of infestation, neonate aphids were gently transferred to 25-day-old Arabidopsis plants, one neonate per plant. To prevent aphid movement between plants, pots were spaced out and placed on the lid of an inverted Petri dish in a tray with soap water. Aphids were checked daily for wing development to prevent dispersal throughout the experiment, but no winged aphids were observed. All plants remained in the vegetative stage during the experiment. The total number of aphids was assessed 14 DPI with Mann Whitney U tests and displayed with emmeans (Lenth *et al*., 2025) and tidyverse (Wickham *et al*., 2019) in R Software v4.4.1 (R Core Team, 2024).

### Aphid feeding behaviour

*Myzus persicae* feeding behaviour was studied on four-to-five-week-old Arabidopsis plants with electrical penetration graph (EPG) recording (Tjallingii, 1988). One day before the experiment, apterous adult aphids were collected from the rearing and transferred to a Col-0 Arabidopsis plant to habituate to the host species. On the day of the recording, a thin gold wire (18 µm diameter, 1.5 to 2 cm length) was gently attached to the aphid dorsum with water-based silver glue. Each wired aphid was placed on a unique plant and connected via a copper nail to a direct current (DC) electrical circuit connected to a Giga-8 amplifier (http://www.epgsystems.eu). Two or three Giga-8 systems were used during each recording day, ranging from 8 to 24 aphids that were simultaneously recorded. Both wild-type and mutant plant lines were included and alternated between channels on each recording day. Electrical signals associated with stylet activities were recorded for eight hours. Waveforms were annotated with EPG Stylet software (http://www.epgsystems.eu) and further processed in R (R Core Team, 2017) with slight adaptations from Kloth et al. (2021). Waveforms that did not occur were considered missing values when calculating the mean and maximum duration and treated as the end time of the recording for latency variables. Events that were interrupted by the end of the recording were not excluded from calculations. R Software v4.4.1 (R Core Team, 2024) was used for further data analysis and visualization. For the summary variables in Supplementary Table S4, χ^2^ tests were performed on proportional data and Mann–Whitney U tests on other variables. To assess temporal patterns in aphid behaviour, the relative proportion of time spent phloem feeding per hour was tested. As these data consisted of many zero’s (hours when aphids did not perform phloem feeding), they were analysed with zero-inflated regression models with time (hour), plant line and their interaction as fixed factors, and aphid individual as random factor, using the zeroinfl function of the “pscl” package (Zeileis *et al*., 2008) in R Software v4.4.1 (R Core Team, 2024).

### Nematode hatching

*Heterodera schachtii* cysts (Woensdrecht population from IRS, Dinteloord, the Netherlands) were collected from sand of infected *Brassica oleracea* plants as previously described (Baum *et al*., 2000). *Heterodera schachtii* were rinsed and transferred into a clean Erlenmeyer. Water was added to a maximum volume of 100 mL containing 0.02% sodium azide. This mixture was stirred for 20 min. Later, sodium azide was vigorously removed by washing with tap water. Cysts were then placed on a hatching sieve in a glass Petri dish. An antibiotic solution was added containing 1.5 mg/mL gentamycin, 0.05 mg/mL nystatin and 3 mM zinc chloride. The cysts were incubated in the dark for four to seven days. Eventually, nematode juveniles were collected in a 2 mL Eppendorf tube. The juvenile stage two (J2) nematodes were surface sterilized with a mercury(II) chloride solution (0.002% Triton X-100 w/v, 0.004% sodium azide w/v, 0.004% mercury(II) chloride w/v) for 20 min. After incubation, the nematodes were spun down and the supernatant was removed. Nematodes were washed with sterile water and spun down again. This was repeated three times. Finally, the nematodes were resuspended in 0.7% gelrite (Duchefa Biochemie, Haarlem, the Netherlands).

### Nematode infection assay

Arabidopsis seeds were vapor sterilized and stratified as described (Willig *et al*., 2024). For *in vitro* nematode assays, seeds were sown in 12-well plates containing modified Knop medium (Sijmons *et al*., 1991). 14-day-old seedlings in a 12-well plate were inoculated with ∼250 sterile J2 *H. schachtii* juveniles in 5 µL gelrite (Baum *et al*., 2000). At 14 and 28 DPI, the number of J3 stage nematodes, males and female nematodes were counted. The counting and imaging were done under a light microscope (ZEISS microscopes and cameras, ZEN 3.2 (Blue edition)). ImageJ (1.53c) was used to measure the size of the females and their corresponding feeding sites. The longitudinal expansion was measured by taking the absolute length of the syncytium. Radial expansion was measured at the widest point of the syncytium. The infection assay was repeated three times independently. The statistical significance of the plant genotypes was assessed with a One-way ANOVA with a post-hoc Tukey HSD test after pooling the data. Data analysis and visualization was performed with R Software v3.6.3 (R Core Team, 2019) using the packages “tidyverse”, “ARTool”, and “multcompView” (Wickham *et al*., 2019; Elkin *et al*., 2021; Graves *et al*., 2024).

### Localization of SLI1 during nematode infection

Arabidopsis *pSLI1:EYFP:SLI1* seeds were sown and grown vertically on a 15cm-by-15cm square Petri dish containing modified Knop medium, at 26°C. Four-day-old seedlings were inoculated with ∼10 sterile *H. schachtii* J2s. At 21 DPI, roots that had a well-established syncytium were selected using an Olympus SZX10 binocular. Thereafter, roots with well-established syncytia were placed on an objective glass with double-sided tape used as spacers and imaged with a ZEISS LSM 510 confocal laser microscope. *pSLI1:EYFP:SLI1* seedlings were excited at 514 nm, and the emitted wavelength was detected on a 517-598 nm emission filter.

## Results

### SLI1 does not affect systemic potyvirus transport

To investigate whether SLI1 affects systemic transport of TEV, two independent *Arabidopsis* T-DNA insertion lines were compared to a positive control, the T-DNA insertion line *rtm2-1* that allows for systemic TEV transport, and their wild-type background line Col-0. Rosette leaves of 14-day-old plants were mechanically inoculated with fresh TEV inoculum, followed by RT-qPCR analysis on the inflorescence at 17 DPI (Fig. 1A). TEV was not detected in the systemic inflorescence tissues of the T-DNA insertion lines *sli1-1* and *sli1-2*, and the wild-type Col-0 (Fig. 1B). In some cases, average Cq values above 34 were observed in *sli1-1, sli1-2*, and Col-0 samples, but aberrant melt curves indicated the absence of TEV in these inflorescences. In contrast to the wild-type Col-0, inflorescence tissues of *rtm2-1* plants tested positive for TEV, giving average Cq values below 32 in nine out of ten replicates. This outcome validates that the mechanical inoculation was successful and confirms the restrictive role of RTM2 in TEV systemic transport. Together, these data indicate that SLI1 does not impair TEV systemic transport.

**Figure 1.**
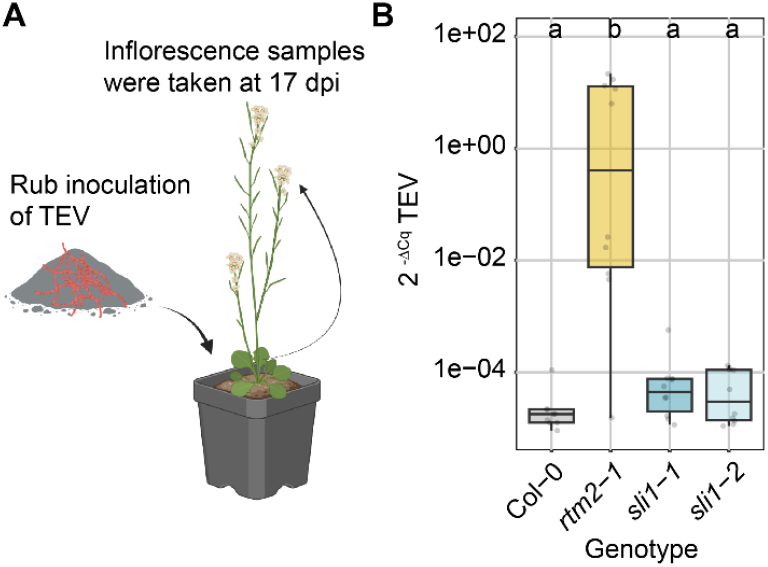
Relative TEV titres at 17 DPI in systemic inflorescence tissues of *Arabidopsis* wild-type Col-0, and the T-DNA insertion lines *rtm2-1, sli1-1*, and *sli1-2*. **(A)** Illustration of how the experiment was carried out. Fourteen-day-old plants were rub-inoculated with TEV, and inflorescence samples were taken 17 DPI. **(B)** Virus titres were determined by RT-qPCR and presented as normalized and linearized relative expression (2^-ΔCq^), displayed on a logarithmic scale. Significant differences between the different plant lines are indicated by different letters above the boxes, as determined by the Kruskal-Wallis test (corrected Dunn) and the Benjamini-Hochberg method to correct for multiple testing (n = 10, p ≤ 0.0022). Boxplots show the median (line), interquartile range (box), extremes (whiskers), and individual data points (dots). Schematic illustration was created in https://BioRender.com.

### Only RTM3 affects aphid reproduction but not phloem feeding

To study the role of RTM in aphid-plant interactions, for each *RTM* gene (*RTM1, RTM2*, and *RTM3*), two independent Arabidopsis T-DNA insertion lines and their wild-type background lines were tested for *M. persicae* aphid feeding behaviour in the first eight hours of infestation and reproduction over the course of two weeks. Previous studies already demonstrated loss-of-function of *RTM2* in *rtm2-1* (Decroocq *et al*., 2006). Other T-DNA insertion lines were confirmed to have significant downregulation of the gene of interest, except for non-significant trends of overexpression in *rtm3-4* (P=0.06, Mann Whitney U test), and downregulation in *rtm2-4* (P=0.10, Mann Whitney U test, Supplementary Fig. S1, Table S2). As it is challenging to obtain accurate transcriptional data of genes with expression limited to immature sieve elements, we decided to continue the bioassays with all T-DNA lines. We observed subtle effects of RTM1 and RTM2 on aphid feeding behaviour (Fig. 2A and B), but without consent between the two T-DNA insertion lines for the same gene. On *rtm1-2*, for example, we observed a reduced proportion of salivation during the phloem phase and a higher mean duration of phloem feeding, indicative of higher susceptibility (Supplementary Table S4). In contrast, on the second mutant, *rtm1-3*, we observed a significant decrease in time allocated to phloem feeding per hour (Fig. 2A, Supplementary Table S5), but without implications for aphid reproduction on either of the *rtm1* mutants (Fig. 2D). On *rtm2-4*, aphids spent less time phloem feeding and initiated less sustained feeding events compared to the wild-type Col-3 (Supplementary Table S4), suggesting the mutant was more resistant. No differences in feeding were, however, observed between *rtm2-1* and Col-0, nor were there any effects on aphid reproduction in either of the *rtm2* mutants (Fig. 2E). Interestingly, only RTM3 had consistent effects in both T-DNA insertion lines. Aphid reproduction was 20 to 30% higher on both *rtm3* mutants compared to the wild-type Col-0 (Fig. 2F), a result that was confirmed in one to two independent repetitions of the experiment. However, feeding behaviour was not affected (Fig. 2C). Only a reduction in short probes was seen on *rtm3-3* (Supplementary Table S4), but no alterations in activities in the phloem, where RTM3 localizes (Sanden and Schulz, 2022). Overall, these findings indicate no major effects of RTM1 and RTM2, but decreased aphid reproduction by RTM3 without obvious effects on feeding behaviour.

**Figure 2.**
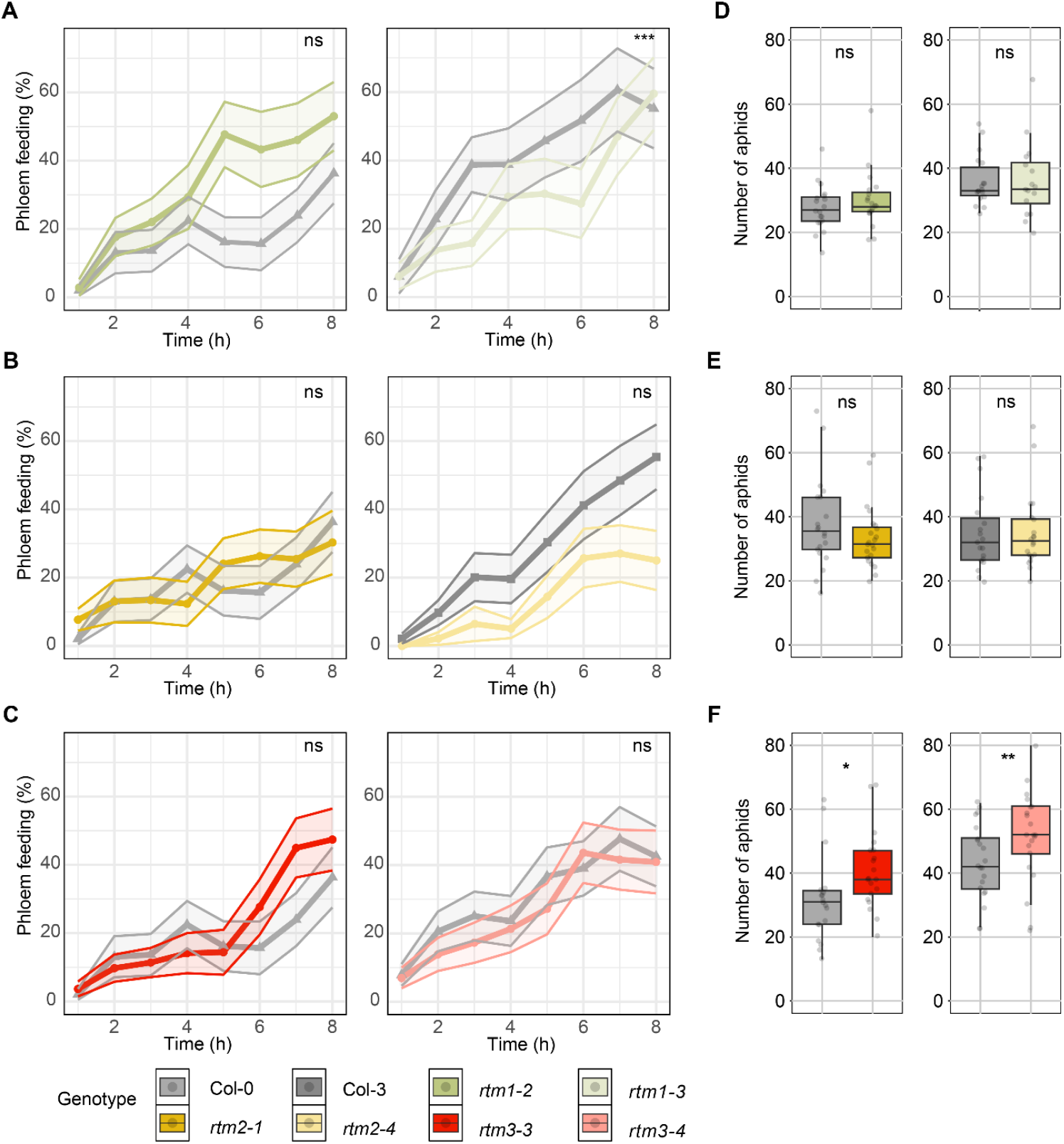
*Myzus persicae* phloem-feeding duration and reproduction on two independent *Arabidopsis* T-DNA insertion lines for *RTM1, RTM2*, and *RTM3*. **(A, B, C)** Phloem feeding on four-to-five-week old Arabidopsis was recorded by electrical penetration graphing (EPG) over eight hours (Col-0/*rtm1-2*/*rtm2-*1/*rtm3-3*: n= 21; Col-0/*rtm1-3*: n= 16; Col-3/*rtm2-4*: n= 20; Col-0/*rtm3-4*: n= 24-25). **(D, E, F)** The number of aphids was recorded 14 days after inoculating individual plants with one neonate aphid (Col-0/*rtm1-2*: n= 19; Col-0/*rtm1-3*: n=16-18; Col-0/*rtm2-1*: n= 20-22; Col-3/*rtm2-4*: n= 19-20; Col-0/*rtm3-3*: n= 19-20; Col-0/*rtm3-4*: n= 21-25). Significant differences in the number of aphids were determined with Mann Whitney U tests, and for phloem feeding with a zero-inflated regression model with Plantline * Time as fixed factors and aphid individual as random factor (ns = not significant, * = p ≤ 0.05, ** = p ≤ 0.01, *** = p ≤ 0.001). Boxplots show the median (line), interquartile range (box), extremes (whiskers), and individual data points (dots), line plots show mean values (dots) with SE in shaded area.

### SLI1, RTM2, and RTM3 support infection and growth of cyst nematodes

To assess whether SLI1 plays a role in cyst nematode infection, we monitored juvenile development and determination of sex of *H. schachtii* in roots of *Arabidopsis* T-DNA insertion lines *sli1-1* and *sli1-2* for four weeks after inoculation. At 14 DPI (Fig. 3A), significantly fewer nematodes developed from infective J2s into third stage juveniles (J3s) on *Arabidopsis sli1-1* and *sli1-2* compared to the wild-type Col-0. At 28 DPI, the total number of nematodes per plant was still significantly less in *sli1-1* and *sli1-2* compared to the wild-type Col-0, pointing at a decline in susceptibility (Fig. 3B). Furthermore, poorer nutritional conditions for *H. schachtii* in Arabidopsis at the early stages of development can lead to the differentiation of more juveniles into males. However, we found that the number of juveniles developing into males and females was still significantly lower in *sli1-1* and *sli1-2* compared to wild-type Col-0 (Fig. 3C and D). At later developmental stages the expansion of syncytial elements limits female growth and thereby offspring size (Urwin *et al*., 1997; Guarneri *et al*., 2024). At 28 DPI, we observed significantly reduced radial expansion of syncytia (i.e., syncytium width) and females growth for the *sli1-1* mutant, but not for the *sli1-2* mutant, compared to wild-type Col-0 (Fig. 3F, H, and I). Nonetheless, the length of syncytia was significantly smaller in both *sli*1-1 and *sli*1-2 mutants (Fig. 3E and G), indicating the incorporation of fewer vascular parenchyma cells. Thus, SLI1 is involved in the establishment of infections by *H. schachtii* during the onset of parasitism, while further on it limits the expansion of syncytia necessary to sustain normal female growth.

**Figure 3.**
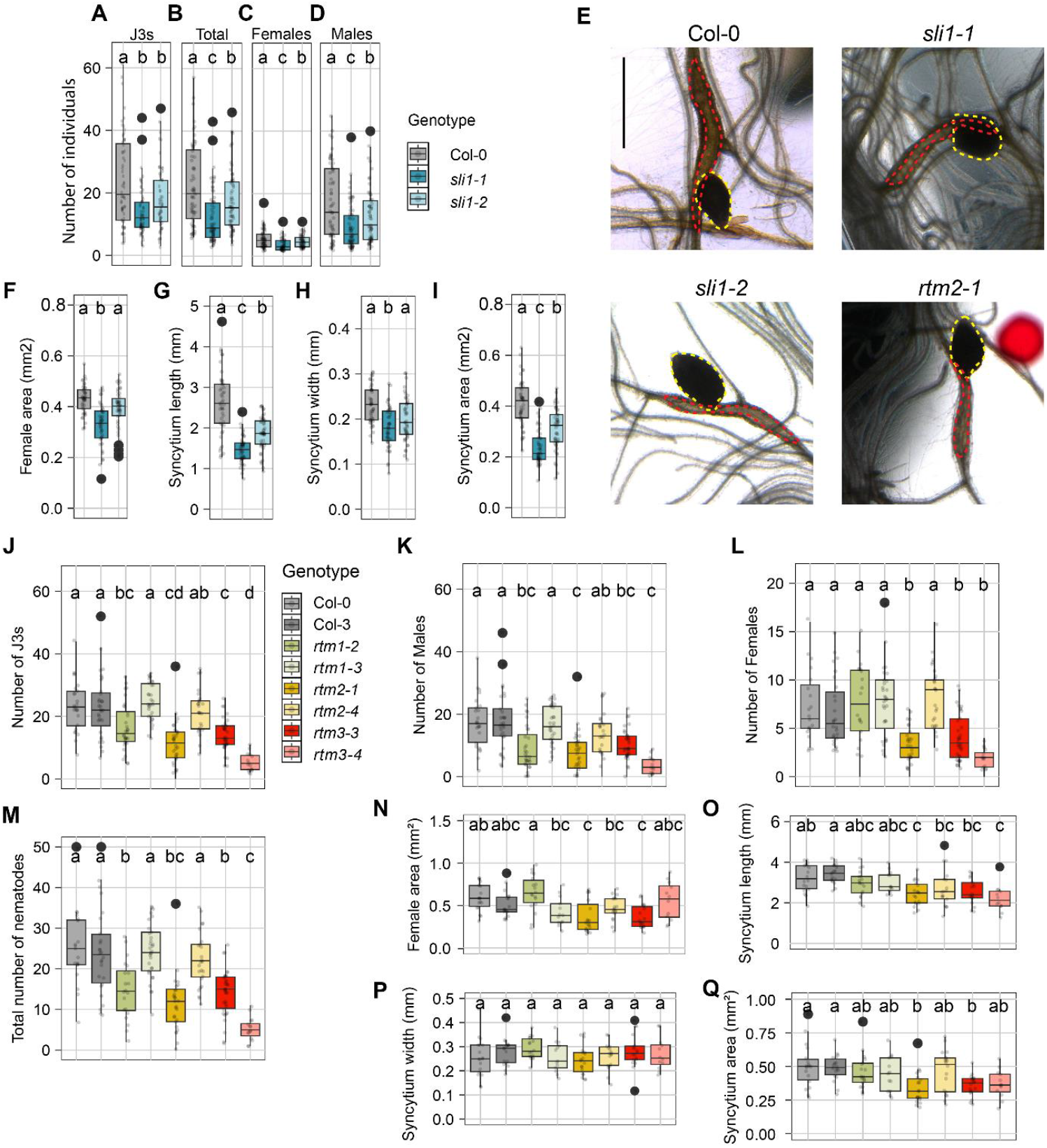
Stationary phloem proteins SLI1, RTM2 and RTM3 are involved in *H. schachtii* infections. Fourteen-day-old Arabidopsis seedlings were inoculated with ∼250 infective juveniles of *H. schachtii*. **(A-I)** *H. schachtii* in *sli1-1, sli1-2*, and Col-0. **(J-Q)** *H. schachtii* in *rtm1-2, rtm1-3, rtm2-1, rtm2-4, rtm3-3, rtm3-4*, Col-0 and Col-3. **(A and J)** The number of third stage juveniles of *H. schachtii* at 14 DPI. **(B and M)** The total number of nematodes at 28 DPI. **(C and L)** The number females at 28 DPI. **(D and K)** The number of males at 28 DPI. **(E)** Representative images of syncytia and adult females at 28 DPI. Scale bar represents 2 mm. Dashed red line indicates the syncytium, and yellow dashed line indicates the cyst. **(F and N)** The maximum two-dimensional surface area of adult females at 28 DPI **(G and O)** The syncytium length. **(H and P)** syncytium width. **(I and Q)** The maximum two-dimensional surface area of syncytium at 28 DPI. Data were pooled and analysed with One-way ANOVA with a post-hoc Tukey HSD test. Letters indicate level of significance (*n* = 25-36 for infection counting; *n* = 12–18 for infection measurements). Boxplots show the median (line), interquartile range (box), extremes (whiskers), outliers (black dots), and individual data points (grey dots).

Next, we assessed whether RTM1, RTM2, and RTM3 influence *H. schachtii* infection using Arabidopsis T-DNA insertion lines *rtm1-2, rtm1-3, rtm2-1, rtm2-4, rtm3-3*, and *rtm3-4*. In case for *rtm3* mutants, a consistent pattern was observed. Both *rtm3-3* and *rtm3-4* consistently showed reduced susceptibility, with significantly fewer J3s, males, and females compared to Col-0. Notably, *rtm3-4* plants gave a stronger phenotype than *rtm3-3* plants. Although minor inconsistencies were observed between the two lines for female size and syncytium dimensions, the overall trend supported a role for RTM3 in promoting infection and syncytium expansion. Results for *rtm1* mutants were inconsistent. Plants carrying the *rtm1-2* mutation showed a significant reduction in the number of J3s, males, and total infections compared to Col-0, but female number, size, and syncytium measurements were unaffected (Fig. 3J–Q). In contrast, *rtm1-3* mutants were not different from Col-0 across all measurements. An inconsistent pattern was observed for *rtm2* mutant plants. Plants carrying the *rtm2-1* mutation showed a clear reduction in nematode performance, with significantly fewer J3s, males, females, smaller female size, and reduced syncytium length and area. However, *rtm2-4*, the only mutant with a Col-3 background and non-significant downregulation of *RTM2*, behaved similarly to the wild type, showing no significant differences. In summary, these findings indicate that the stationary phloem protein RTM1 has no or marginal effects on *H. schachtii*, but that SLI1, RTM3, and possibly RTM2 support cyst nematode infection, female growth, and expansion of syncytial elements.

### SLI1 localises near nematode-induced syncytia and adjacent sieve tubes

Cyst nematodes induce *de novo* formation of sieve tubes, in which SLI1 is normally localized in low abundance (Kloth *et al*., 2017). To determine if SLI1 localizes around the syncytium or in newly formed sieve tubes, we examined the localization of SLI1 in *H. schachtii-*infected roots of Arabidopsis using a transgenic Arabidopsis line expressing SLI1 fused to YFP under its native promotor (*pSLI1:EYFP:SLI1*). Fluorescent imaging revealed brighter fluorescent signals around infection sites of *H. schachtii* compared to the signal observed in non-infected tissue (Fig. 4). Moreover, the diameter of SEs around syncytia increased compared to uninfected roots, indicating a high abundance of SLI1 proteins in the direct surrounding of syncytia.

**Figure 4.**
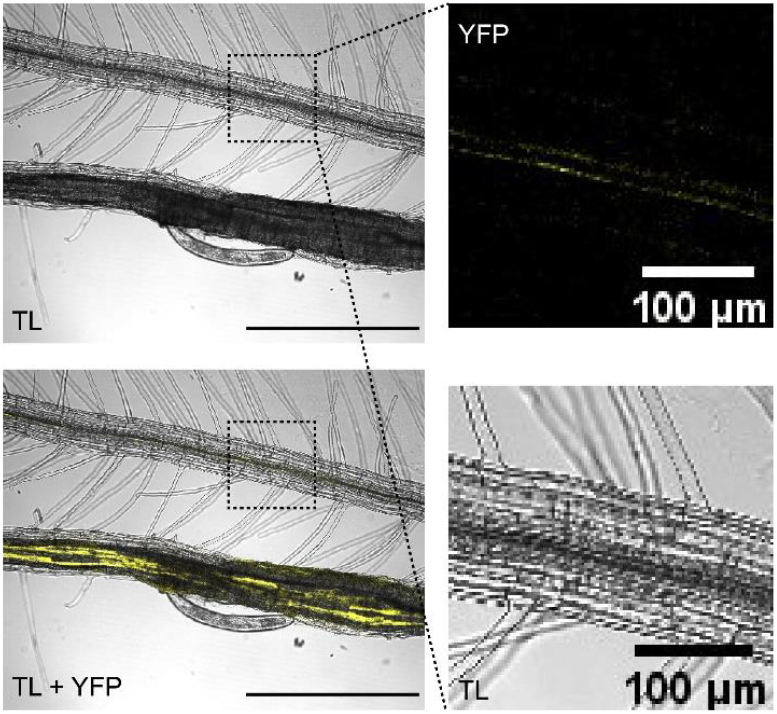
SLI1 accumulates in *de novo* formed sieve tubes. Seven-day-old Arabidopsis seedlings expressing *pSLI1:EYFP:SLI1* were inoculated with ∼10 *H. schachtii* juveniles at 26°C. Fluorescent signals were imaged at 21 DPI. Confocal images of *pSLI1:EYFP:SLI1* (yellow) in roots with details in the boxed region showing pSLI1:EYFP:SLI1 in uninfected tissue. The experiment has been repeated three separate times and similar phenotypes were observed (n = 3). TL is transmission light. YFP is fluorescent signal of *pSLI1:EYFP:SLI1*.

## Discussion

Various pathogens, such as plant viruses, phloem-feeding insects, and cyst nematodes, completely depend on the phloem for their systemic transport, feeding, and successful reproduction, respectively. To counteract these intruders, plants deploy various defence mechanisms in different tissues or cell types, but our understanding of phloem-based resistance is limited. Previously, the phloem proteins RTM1, RTM2, and RTM3 were shown to restrict long-distance transport of potyviruses (Mahajan *et al*., 1998; Whitham *et al*., 2000; Chisholm *et al*., 2001; Cosson *et al*., 2010), while SLI1 confers broad-spectrum resistance to phloem-feeding insects, including the green peach aphid (Kloth *et al*., 2021). However, the broad-spectrum effects of RTMs and SLI1 on potyviruses and aphid remained unknown, and the role of both proteins in cyst nematodes infections had not yet been explored. To this end, this work aimed to elucidate the spectrum of resistance that is conferred by the phloem proteins SLI1, RTM1, RTM2, and RTM3 against TEV (potyvirus), *M. persicae* (aphid), and *H. schachtii* (cyst nematode). We found contrasting findings for the function of SLI1 and RTMs in plant-virus, -aphid, and -nematode interactions.

### Resistance specificity of SLI1 and the RTMs

*SLI1* and *RTM2* both encode transmembrane-domain containing hsp20-like proteins that are limited to phloem SEs (Chisholm *et al*., 2001; Cayla *et al*., 2015; Kloth *et al*., 2017, 2021). Even though both proteins belong to the same protein family, this study demonstrates that they do not share the same functionality. SLI1 is unrelated to the restriction of TEV systemic transport, as loss-of-function *sli1* mutants did not support TEV systemic transport and showed similar virus systemic movement as the wild type Col-0. Additionally, no consistent differences were found in *M. persicae* phloem-feeding behaviour or reproduction between the loss-of-function *rtm2* mutants and wild type line. Therefore, even though SLI1 and RTM2 are closely related hsp20-like proteins (Bondino *et al*., 2012) that both reside in the sieve tubes, our results indicate that their function is not redundant but specialized to confer resistance to either *M. persicae* or TEV, respectively. Furthermore, it is likely that these pathogen-specific resistances are mediated by different mechanisms. For instance, SLI1 might be involved in an occlusion mechanism (Kloth *et al*., 2021), while RTM2 might interact with other (RTM) proteins to interfere with TEV systemic transport by sequestering virus proteins or movement complexes. Possible hypotheses on the underlying mechanism facilitating SLI1- and RTM2-mediated resistance are extensively discussed in the review by Kloth & Kormelink (2020).

Previously, RTM3 was demonstrated to restrict potyvirus systemic transport (Cosson *et al*., 2010), and in this work it was the only protein that also reduced *M. persicae* reproduction, as loss-of-function mutants supported a ∼30% increase in aphid reproduction. However, the effect of *rtm3* on aphid reproduction is less prominent compared to the ∼50% reproduction increase observed in *sli1* loss-of-function mutants (Kloth *et al*., 2017). In addition, we did not observe an increase in phloem-feeding as seen in *sli1* loss-of-function mutants, and therefore consider RTM3 to be involved in a pathway distinct from SLI1-mediated resistance. It is possible that the effects of RTM3 are induced over a longer time frame or accumulate more slowly in the aphid and therefore remained undetected within the eight hours of EPG monitoring. While most phloem-located defences in Arabidopsis seem to act within minutes or hours, such as SLI1-, PAD4- and EDR1-like-mediated mechanisms (Pegadaraju *et al*., 2007; Kloth *et al*., 2017; Guo *et al*., 2020), the OPDA-regulated lipase MPL1 is another example of a phloem-associated protein involved in successful suppression of aphid reproduction without obvious effects on *M. persicae* feeding in the first eight hours of contact (Louis *et al*., 2010; Archer *et al*., 2023). This illustrates that some phloem-associated defences require more time, perhaps one or more days, before they affect aphids. Presumably, the latency to a defence phenotype may inform us about the complexity of the involved metabolic pathways or gradual accumulation of toxic products in the aphid body.

### *SLI1, RTM2, and RTM3* are involved in cyst nematode infection

The cyst nematode *H. schachtii* relies on the phloem for nutrients to support its development and reproduction, making it likely to be affected by phloem-based resistance mechanisms. Loss-of-function mutants of *sli1, rtm2*, and *rtm3* reduce Arabidopsis susceptibility to *H*. schachtii by reducing the numbers of J3 juveniles, males, and females. Next, for *sli1-1*, we observed that syncytia were significantly smaller due to reduced longitudinal and radial expansion of the feeding structure. For *rtm2* and *rtm3* mutants we found that syncytia were also significantly smaller, but this could only be attributed to reduced longitudinal expansion. Together, these findings suggest that Arabidopsis *SLI1, RTM2, and RTM3* are susceptibility genes in the case of *H. schachtii* infection. Unlike their protective role against *M. persicae* and TEV, *SLI1* and the *RTMs* apparently have counteractive effects on plant protection against cyst nematodes. Similarly to the aphid assays, RTM1 appears to have little or no effect on nematode reproduction, indicating some level of specificity among phloem proteins in their role in supporting cyst nematode infection.

The effect of RTM2 on nematode reproduction is not consistent between the two individual loss-of-function mutants *rtm2-1* and *rtm2-4*. Further validation of *rtm2-1* with a second infection assay (Supplementary Fig. S2) gave similar outcomes as we first observed (Fig. 3). It is possible that *rtm2-4* is not a complete RTM2 loss-of-function mutant (Whitham *et al*., 2000) as is illustrated by the non-significant downregulation of RTM2 by the insert in the promotor region (Supplementary Fig. S1). The expression level of *RTM2* in the *rtm2-4* mutant could have been sufficient to support cyst nematode infection. The phenotype of *rtm2-1* is more reliable, as Decroocq *et al*. (2006) already proved loss-of-function of the resistance to plum pox virus in this mutant line.

### Phloem proteins modulate syncytium expansion

In contrast to potyviruses and aphids, cyst nematodes require extensive morphological changes in host cells to form a syncytium, which serves as their primary feeding site. This structure provides access to the phloem, ensuring a steady supply of nutrients essential for nematode development. We believe that SLI1, RTM2, and RTM3 promote longitudinal syncytium expansion, supporting female nematode growth and likely also fecundity. In *rtm2-1, rtm3-3*, and *rtm3-4* mutants, syncytia exhibited reduced longitudinal expansion, leading to smaller overall syncytia, while radial expansion remained unchanged. In Col-0 background, we observed that SLI1-eYFP signal was strongest in the longitudinal ends of the syncytium, the site which expands by hypertrophy after cells are no longer incorporated, which occurs approximately after seven DPI (Amjad Ali et al., 2014; Hewezi et al., 2014; Wieczorek et al., 2006).

Interestingly, SLI1 expression is higher in pre-mature SEs compared to mature SEs. Therefore, the lower SLI1-eYFP signal in the centre of the syncytium is probably due to the maturation of SEs, whereas at the ends, premature SEs are still found, and enucleation has not yet occurred. This is in line with earlier findings where companion cells are formed after *de novo* vascularisation, and therefore more CCs found in the centre of the syncytium (Hoth *et al*., 2005; Kim *et al*., 2020). One hypothesis is that SLI1 facilitates longitudinal syncytium expansion by promoting *de novo* SE formation in the surrounding vasculature, thereby improving nutrient delivery and reducing mechanical constraints at the expanding edges of the syncytium.

Given that SLI1, RTM2, and RTM3 all similarly affect *H. schachtii*, they may play a crucial role in vascular remodelling during syncytium formation. Given that these proteins influence longitudinal expansion, they may facilitate *de novo* SE formation, thereby enhancing nutrient flow and turgor driven expansion of the syncytium. Disrupting any of these genes could interfere with the establishment of new phloem connections, ultimately limiting the nematode’s ability to withdraw nutrients from the host. If SLI1, RTM2, and RTM3 are indeed involved in *de novo* vascularization around syncytia, we expect to observe reduced or disrupted vascularization surrounding infection sites in mutant plants. Additionally, live imaging of phloem transport using fluorescent tracers could provide further insights into how these proteins influence nutrient flow into the syncytium. By elucidating their precise role in vascular remodelling, we may better understand how nematodes manipulate host phloem to establish, expand, and sustain their feeding structures.

### Balancing resistance and susceptibility

While reducing the proliferation of a specific pathogen may seem beneficial, it can lead to unintended increases in others, highlighting the complexity of plant defence strategies. Our findings show that while SLI1 and RTM proteins respectively limit aphid and potyvirus multiplication, they also allow greater proliferation of the cyst nematode *H. schachtii*. This suggests that these proteins might have dual effects on pathogen success, possibly through their involvement in vascular remodelling. Such trade-offs may result from resource reallocation, cross-talk between defence pathways, or the ability of certain pathogens to exploit host processes originally aimed at limiting other pathogens. These results underscore the importance of understanding the interconnected nature of plant defence mechanisms before enhancing protection against one threat, to avoid inadvertently improving conditions for another.

Moving forward, future research should focus on uncovering the molecular mechanisms that drive these pathogen-proliferation dynamics. A key approach is to investigate protein-protein interactions involving SLI1 and the RTMs in the Arabidopsis phloem, using techniques such as proximity labelling. Identifying interaction partners will help determine whether these proteins primarily function in signalling, phloem transport, or vasculature formation. Additionally, studying their effects on SE function at the cellular level will provide further insights into their role in pathogen defence. Beyond individual molecular interactions, a broader systems biology approach including transcriptomic and metabolomic analysis could help identify additional factors influencing phloem-based defences. Combining these methods will allow for a more comprehensive understanding of plant defence mechanisms and aid in developing strategies that reduce multiple pest and pathogen burdens simultaneously.

## Abbreviations

DPI: Days post-inoculation
EPG: Electrical penetration graph
J2: Juvenile stage two
J3: Juvenile stage three
RTM: RESTRICTED TEV MOVEMENT
SE: Sieve element
SLI1: SIEVE ELEMENT-LINING CHAPERONE1
TEV: Tobacco etch virus

## Supplementary data

The following supplementary data are available at JXB online.

Table S1. PCR primer details for selection of homozygous T-DNA lines.

Table S2. RT-qPCR primer details for assessment of transcription of the gene of interest in T-DNA lines.

Table S3. RT-qPCR primer details used for the detection of TEV.

Table S4. *Myzus persicae* aphid probing behaviour during 8-hr electrical penetration graph recordings.

Table S5. Effect of plant line on the proportion of time spent phloem feeding per hour.

Fig. S1. Relative expression for assessment of transcription of the gene of interest in T-DNA lines.

Fig. S2. Repetition of nematode infection assay as validation of *rtm2-1*.

## Acknowledgements

We would like to acknowledge Biointeractions & Plant Health of Wageningen Research for kindly providing the TEV strain, the insect rearing team at the Laboratory of Entomology at Wageningen University & Research for maintaining the *Myzus persicae* rearing, and Tamara Kalsbeek for her contributions to the EPG recordings.

## Author contributions

JD, KJK, EEM, and JJW: Conceptualization; JD, KJK, EEM, and JJW: methodology; NC, JD, JH, JNK, KJK, and JJW: investigation; KJK, and JJW: supervision; JD, KJK, and JJW: visualization; JD, KJK, and JJW: writing – original draft preparation; JD, MD, KJK, EEM, MMO, GS, and JJW: writing – review & editing

## Data availability

Raw data collected in this study is available upon request.

